# Multimodal generation of astrocyte by integrating single-cell multi-omics data via deep learning

**DOI:** 10.1101/2023.11.30.569500

**Authors:** Jiashun Mao, Jianmin Wang, Amir Zeb, Kyoung Tai No

## Abstract

Obtaining positive and negative samples to examining several multifaceted brain diseases in clinical trials face significant challenges. We propose an innovative approach known as Adaptive Conditional Graph Diffusion Convolution (ACGDC) model. This model is tailored for the fusion of single cell multi-omics data and the creation of novel samples. ACGDC customizes a new array of edge relationship categories to merge single cell sequencing data and pertinent meta-information gleaned from annotations. Afterward, it employs network node properties and neighborhood topological connections to reconstruct the relationship between edges and their properties among nodes. Ultimately, it generates novel single-cell samples via inverse sampling within the framework of conditional diffusion model. To evaluate the credibility of the single cell samples generated through the new sampling approach, we conducted a comprehensive assessment. This assessment included comparisons between the generated samples and real samples across several criteria, including sample distribution space, enrichment analyses (GO term, KEGG term), clustering, and cell subtype classification, thereby allowing us to rigorously validate the quality and reliability of the single-cell samples produced by our novel sample method. The outcomes of our study demonstrated the effectiveness of the proposed method in seamlessly integrating single-cell multi-omics data and generating innovative samples that closely mirrored both the spatial distribution and bioinformatic significance observed in real samples. Thus, we suggest that the generation of these reliable control samples by ACGDC holds substantial promise in advancing precision research on brain diseases. Additionally, it offers a valuable tool for classifying and identifying astrocyte subtypes.

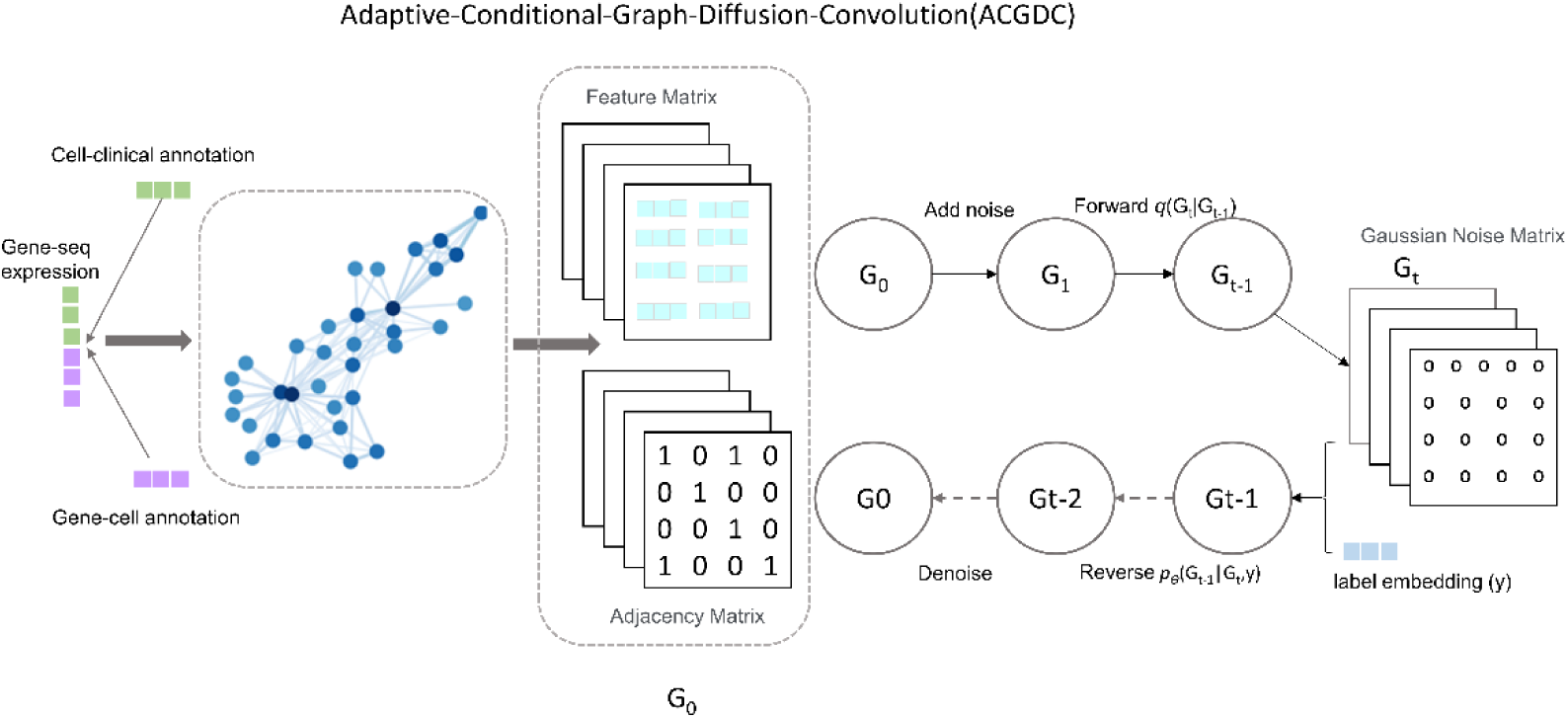

## I. INTRODUCTION

### 1. Background on astrocytes

Glial cells including astrocytes[1], and oligodendrocytes and their precursor cells as well as microglia provide physiological and morphological platforms for the central nervous system (CNS). Among them, Astrocytes play a critical role in neurotransmitter regulation, ion homeostasis, maintenance of the blood-brain barrier, and production of basement membrane and perineural trophic factors [2–6]. In addition, it has been investigated that these astrocytes are incorporated with ion channels such as K+, NA+, CA+ [7–9]. For example, substantial evidence indicates that increased Ca2+ concentrations play an important role in information transmission between astrocytes and between astrocytes and neuronal cells [10]. The biological effects due to changes in Ca2+ concentration in astrocytes mainly include:

1. Intracellular ionic oscillations due to intracellular Ca2+ release,
2. The formation of Ca2+ efflux during neural activity by excitation of transmitters such as glutamates and purines,
3. Induced release of transmitters such as glutamates from astrocytes into the extracellular gap triggering receptors leading to neuronal formation of currents,
4. Transmission of cellular signals to neighboring astrocytes [11–14].

Thus, astrocytes wrap around synapses and play an important role in maintaining their humoral, ionic, acid-base environment and endostasis [15], even guaranteeing normal inter-synaptic information transfer. Normally, astrocytes are categorized into static, activated, and proliferating states, and their dynamic transition between these three states collectively constituting the broader concept of the cell cycle [16].

In the normal central nervous system, both static and activated glial cells co-exist. However, these life-shaping cells respond to injury and disease in pathophysiological conditions [17–19] when exposed to injury, the static state undergoes a gradual transition into the activated state which is leveraged by the influence of cytokines [20–24]. Besides their role in brain tissue injury, for instance, the formation of glial scars after spinal cord injury [25], astrocytes also play a pivotal role in the investigation of neurological diseases like Glioblastoma (GBM) and Alzheimer’s disease (AD) [10, 26, 27]. Additionally, they play an essential role in fundamental biological mechanisms, including neuronal signaling (associated with memory, learning, sleep mechanism[28]), neuronal metabolism, growth, and communication. In contrast to astrocytes in the activated and proliferating states, the evaluation of astrocytes in the static state can also provide valuable insights into regulatory mechanisms of molecules within cells undergoing extraordinary conditions. This is because the static state serves as a reference point for the activated state, thereby aiding in the identification of molecular mechanisms underlying cellular abnormalities. Thus, in turn, it can offer precious ideas for promising treatment avenues.

To this end, data from astrocytes in healthy tissues offer valuable bioinformatics significance, serving as an essential reference group for differential analysis, enrichment analysis, and other research platforms in the exploration of disease mechanisms. However, acquiring such data faces significant challenges due to various reasons including cost and difficulty associated with collection. Moreover, considering the perspective of precision medicine, individual variations withing these samples further complicate the research investigation. Consequently, a substantial and diverse collection of astrocytes samples becomes imperative to comprehensively study the pathology and brain homeostasis mechanisms underlying these brain diseases.

The recently emerged single-cell multimodal omics (scMulti-omics) sequencing technologies have revolutionized our capabilities, allowing for the simultaneous detection of multiple modalities within the same individual cell, such as RNA expression[29], protein omics, and chromatin accessibility [29, 30]. These groundbreaking technologies not only explore the intricate cellular heterogeneity across multiple molecular levels, but also establish meaningful connections across different layers, offering a systematic view of interactions and regulations at the single-cell resolution. By incorporating multiple measured modalities in the analyses of biological samples, these advancements significantly enhance the potential for groundbreaking insights into the molecular mechanisms underlying numerous processes, such as cell physiology, tissue development, and disease prevalence.

The growing size of scMulti-omics datasets necessitates to create novel and efficient computational tools capable of integrating massive high-dimensional data generated from diverse sources. This integration is crucial for enabling more comprehensive and reliable downstream analyses, thus leveraging knowledge mining [31]. Such concept of “integrative analyses” not only substantiate the construction of a large-scale single-cell multimodal atlas, but also fulfills an urgent need to fully utilize publicly accessible single-cell multimodal data [32]. Such an atlas can essentially serve as an encyclopedia, enabling researchers to transfer their knowledge to new datasets and in-house studies [33–35].

On the contrary, several obstacles must be overcome to computationally extract meaningful knowledge from the highly intricate single-cell multimodal data. Firstly, the inference of low-dimensional joint representations from multiple patterns that can be applied in downstream analysis like clustering, is a challenging task. Secondly, while multimodal single-cell data allows for the learning of relationships between patterns that can be used to train predictive models, obtaining high accuracy in many-to-many predictions of single-cell data (for instance from single-cell transcriptomes to chromatin accessibility) remains an unresolved issue. These problems primarily stem from the intricate nature of capturing common underlying factors and relationships across modalities that significantly differ in characteristics, such as data distribution, dimensionality, and sparsity.

### 2. Methods for integrating single-cell multi-omics data

Deep learning offers an extensive array of applications and successful case studies in research disciplines like computational biology and the exploration of novel drug. Significant advancements are being devised in the application of deep generative models across various areas, including molecular design, protein engineering, chemical and biological space exploration, and data augmentation [36]. Subsequently, deep generative models are gaining high fame and demonstrating increasing effectiveness in the generation of single-cell samples [37]. These generative models primarily categorized into two types: one type is based on gene expression data from single-cell transcriptome sequencing, where the genomic expression matrix of each cell serves as input. These models utilize mainstream generative approaches, including scGAN(single-cell Generative Adversarial Network), cscGAN (conditioning single-cell Generative Adversarial Network)[38], Splatter and SUGAR[39, 40], LSH-GAN [41]. These models learn to train an intermediate hidden vector and subsequently reconstruct a new single-cell gene expression matrix to generate novel single-cell samples. The other type of generative model is rooted in the data structure of graph networks, which are employed to establish connections between gene molecules and other gene influencing molecules [42–48]. Such models execute graph neural network algorithms for several training tasks, including inter-molecular relationships, node classification, and even graph generation as seen in GLUE[49]. However, a significant limitation lies in the fact that the majority of these methods are based solely on individual omics data, and their performance tends to be suboptimal when dealing with multi-omics and heterogeneous networks.

Recently, several methods proposed to integrate single-cell multimodal data. While, these methods are designed to tackle the tasks such as extracting latent features, their current performance is limited in different ways. For instance, methods based on generalized linear models like Seurat lib and scAI model [50, 51] often struggle to capture the intricate structures present in single-cell data [37]. On the hand, there are variational autoencoder-based (VAE) methods available, including scMVAE and totalVI, for the analyses of single-cell multimodal data [52, 53]. Nevertheless, scMVAE needs the reduction of chromatin accessibility to the transcriptome prior to training, a step which is known to result in non-negligible loss of epigenetic information [50, 54].

The potential to simultaneously analyze both the transcriptional and chromatin landscapes of single cell has evolved into a powerful technique, enabling the identification of distinct cell population and elucidating insights into the regulation of gene expression within the population. MIDAS is a deep generative model, which is based on VAE architecture[55, 56]. MIDAS is specifically designed to capture the joint distribution of incomplete single-cell multimodal data, incorporating measurements from Assay for Transposase-Accessible Chromatin (ATAC), RNA, and Antibody-Derived Tags (ADT). The input to MIDAS is a mosaic feature-by-cell count matrix that encompasses various single-cell samples (batches). The outputs from this model are suitable for a range of downstream analyses, and cell typing [56, 57].

MultiVI [58] is a probabilistic framework that utilizes deep neural networks to synergistically analyze multimodal single-cell data types, including single cell RNA data (scRNA), single cell transposase-accessible chromatin data (scATAC) and the combination of both (scRNA + scATAC) data. This approach generates a low-dimensional latent space that is rich in information and accurately reflects the chromatin and transcriptional properties of cells, even in the scenario when one of the modalities is missing. MultiVI takes into account technical effects present in scRNA and scATAC-seq data, while correcting batch effects present in both data modalities.

The single-cell Multi-View Profiler (scMVP) [59]is an innovative approach which is capable of simultaneously measuring gene expression and chromatin accessibility within the same individual cell. scMVP employs an automatic learning process to establish a shared latent representation for scRNA-seq and scATAC-seq data. This is accomplished via a multi-view VAE constrained by clustering consistency. scMVP further estimates each layer of data from the multi-omics common latent embedding. It does this through specific data generation mechanisms for each layer, which includes a self-attention-based scATAC generation channel and a scRNA generation channel utilizing mask attention. scMVP serves multiple purposes by creating common latent representations, such as dimensionality reduction, cell clustering, and developmental trajectory inference. It also generates separate imputations for tasks like differential analysis and cis-regulatory element identification. This approach helps mitigate data sparsity concerns through imputation and accurately identifies cell groups, making it ideal for different joint profiling techniques that utilize a common latent embedding.

On contrary, scMM is a novel statistical framework formulated for single-cell multimodal analysis. It leverages datasets generated by advanced technologies like transcriptome and epitope cell index sequencing (CITE-seq) as well as simultaneous high-throughput assays for transposase-accessible chromatin (ATAC) and RNA expression sequencing (SHARE-seq). This framework efficiently extracts multimodal information that encodes biologically significant potential variables [54].

MOFA+ (Multi-Omics Factor Analysis v2) [60] and the WNN dataset [51] have been introduced to address the integration of bimodal data encompassing complete modalities [56]. In contrast, approaches like totalVI [61], sciPENN [62], Cobolt [63], and MultiVI [58] are tailored for bimodal integration where certain modalities might be missing. However, fewer trimodal integration methods are currently accessible for the integration of trimodal data. For instance, MOFA+ has been specifically designed to integrate complete trimodal modalities. Additionally, GLUE [49] and uniPort [64] were developed for the integration of unpaired trimodal data.

The tools elaborated earlier like MultiVI and Cobolt, employ symmetric multimodal Variational Autoencoder (VAE) models for federated modal single-cell datasets. However, in the context of multimodal data integration, the potential embeddings generated by the encoder capture shared semantic features across modalities, while the data generated by the decoder still preserves modality-specific biological information. This specificity is needed to integrate similarities between modalities. When dealing with joint analysis datasets characterized by extreme data sparsity and random noise in either dataset group, the inconsistency of the joint multimodality embedding can substantially distort the biological variation within the cellular latent embedding. This inconsistency could lead to the generated model producing excessively smooth data in the continuous distribution, ultimately, hindering the interpretation and downstream application of the joint latent embedding.

All the existing integration methods are developed to address specific and restricted combinations of omics data. Due to the diversity of scMulti-omics technologies, datasets originating from different studies frequently encompass dissimilar combinations of omics types, which might involve one or more missing modalities, resulting in a mosaic-like data. The prevalence of mosaic-like data is rapidly increasing and can be anticipated to continue growing. Consequently, there is an urgent demand for versatile and comprehensive integration approach to significantly broaden the scope and modalities of integration. Such an integration is necessary to overcome the current limitations in modality scalability and cost associated with currently accessible scMulti-omics sequencing technologies. Nevertheless, multimodal mosaic integration is quite challenging [56, 65]. A key challenge involves addressing the diversity of modalities and handling technical variations across different batches. Another significant challenge is the successful implementation of modality imputation and batch correction, which are crucial for downstream analysis.

To tickle down these challenges, we have leveraged a fundamental framework known as ACGDC (adaptive conditional graph diffusion convolution). This framework is designed to facilitate the mosaic integration and transfer of knowledge within single-cell multimodal data. By harnessing the power of self-supervised learning [66, 67], and information-theoretic approaches [68, 69], ACGDC concurrently achieves modality alignment, imputation, and batch correction for single-cell trimodal mosaic data. Furthermore, we have integrated tailored-transfer learning and reciprocal reference mapping schemes into ACGDC to enable effective knowledge transfer.

Our comprehensive evaluation, involving systematic benchmarks and case studies, underscores that ACGDC can accurately and robustly integrate mosaic datasets. Notably, by applying ACGDC to the atlas-level mosaic integration of multimodal human brain data sourced from The Adult Genotype-Tissue Expression (GTEx) project [70], we have successfully achieved flexible and accurate knowledge transfer across the diverse unimodal and multimodal query datasets.

### 3. The integration method and data source

The integration of single-cell multi-omics data involves a range of diverse methodologies, including data frames [38], and graph networks [71, 72]. Currently, most bioinformatics analyses of sequencing data are built upon the framework of data frames. Nevertheless, such format often contains sparse expression, and may exhibit issues like missing values, abnormal values, and other noisy data, all of which can potentially influence the subsequent analyses. Since, the data expression patterns can differ across different histology, the integration of data from diverse histological sources is often tricky. This challenge is exacerbated when considering additional meta-data, labels, annotations, and patient clinical information. To this end, the adoption of graph networks offers an inherent advantage of circumventing issues of sparsity and heterogeneity. Graph networks encompass both homogeneous and heterogeneous graphs. In this section, our emphasis focuses on the heterogeneous graphs [73, 74].

In this study, we leverage heterogeneous graphs for the representation of single-cell multi-omics. Firstly, we retrieve RNA-Seq expression (gene and transcript level, read counts) and metadata (clinical information) from The GTEx Analysis V8 release which is a comprehensive public resource for the study of tissue-specific gene expression and regulation. All samples were collected from 54 non-diseased tissue sites across nearly 1000 individuals, primarily for molecular assays. Then, we filter out non-brain astrocyte expression data. In addition, we get cell marker data from http://bio-bigdata.hrbmu.edu.cn/CellMarker/index.html for the annotation of cell type according to gene symbol. All the original data can be available at https://doi.org/10.5281/zenodo.8358553.

We integrate message passing and graph attention mechanisms in an unsupervised pre-training phase establish a generic representation for subsequent tasks. Such tasks encompass a variety of applications, including cell subtype classification, cell sample generation, cell state trajectory inference, disease mechanism exploration, identification of therapeutic targets, and discovery of biomarkers. Subsequently, we execute supervised fine-tuning, which is based on specific downstream tasks like cell subtype identification and sample generation.

### 4. Definition of graphs and construction of data sets

#### Definition of graphs

A graph is defined as “a structure to represent entities and their relationships, denoted as G= (V, E)”. Such graph is comprised of two sets: the set V containing nodes and the set E containing edges. Within the set of edges E, an edge (u, v) connects a pair of nodes u and v, signifying a relationship between them. Such a relationship can either be undirected, indicating a symmetric relationship between nodes, or directed, describing an asymmetric link. Graph can be weighted or unweighted. In weighted graphs, each edge carries a scalar weight value. This weight could indicate factors like length or strength of the connection.

Graphs can be divided in two main types: homogeneous or heterogeneous. Homogeneous graphs are comprised of nodes and edges that represent identical entity and relationship types. For instance, in a social network graph, node represents individuals, and edges indicate social connections between them. In contrast, heterogeneous graphs encompass diverse nodes and edge types, often with unique properties. These attributes aim to characterize each node and edge type. In complex scenarios, graph neural networks might require distinct dimensional representations to model different node and edge types.

In this study, we study heterogeneous graphs, for which we define five edge types: hierarchical relations, interaction relations, multimodal relations, composition relations, and sequence relations (as shown in Figure 1). Each node has properties sourced from ontology or statistics, categorized as instance entity (such as molecular components like a RNA, a protein, a DNA, obtain the property from itself) and notion entity (cell class or RNA type with properties derived from calculated statistical values). Every edge possesses its own type and weight, determined by the nodes on both ends. Message passing within the heterogeneous graph can be divided into two main steps: 1) computing and aggregating messages for each adjacent edge; 2) For every node, the new representation is updated from aggregating different relationship edge with different weight.

**Figure 1.**
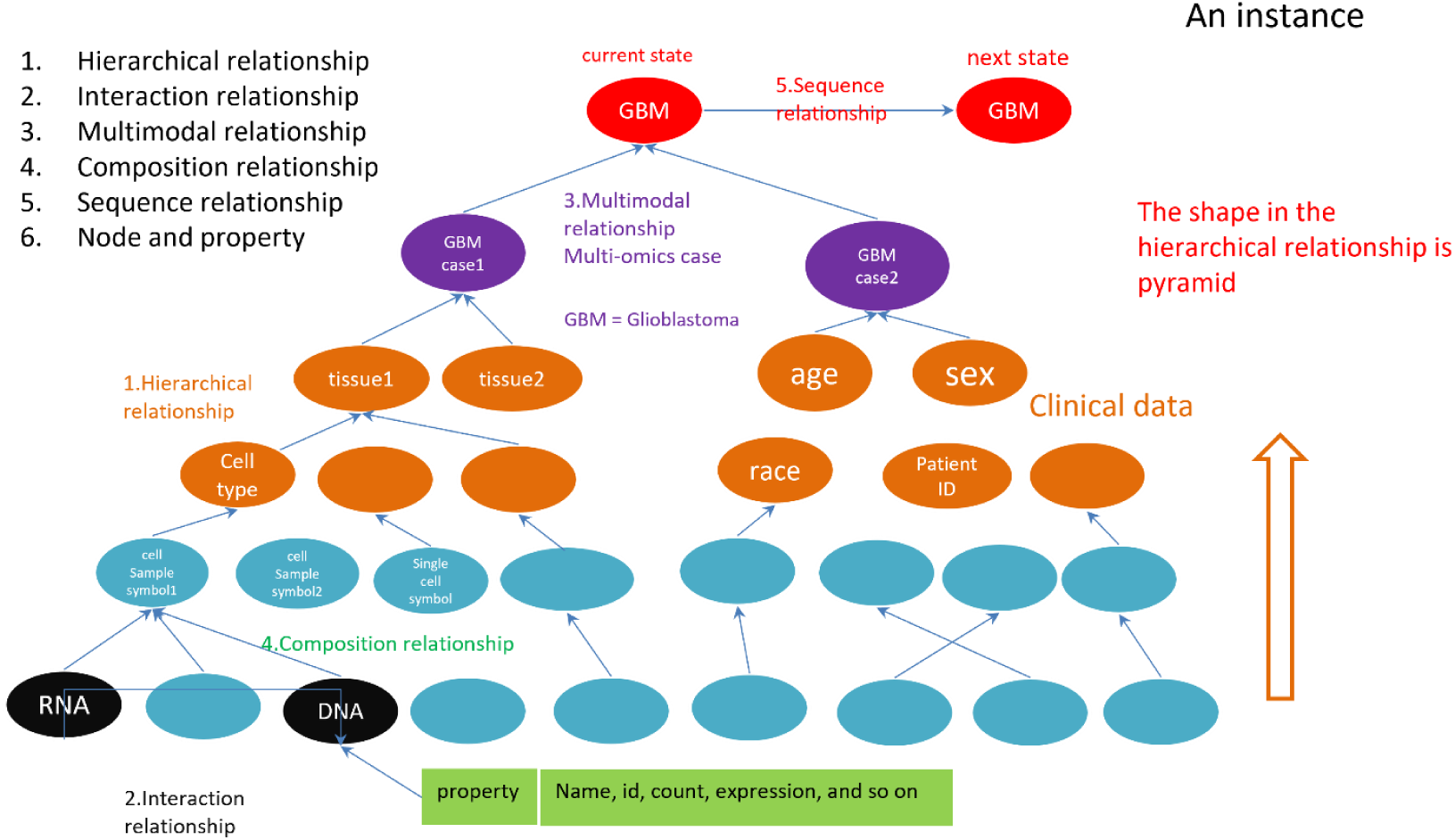
The definition of node relationships and node attributes in our graph network for the integration of single-cell multi-omics data.

#### Construction of data sets

We initiated the process by acquiring sequencing data from the GTEx database, focusing brain tissues and astrocyte. Subsequently, we performed the following procedure on this dataset. Following annotation steps 1) to 6), S1-S4 in the supporting file provides the detail of preprocessing these data. S5-S6 in the supporting file present the statistical description of node types and edge types along with node entity. The outcome of step 7) is illustrated in section 3.1.

1. Id exchange (ensembl_gene_id-> entrezgene_id, hgnc_symbol)
2. Gene/RNA category identification
3. Cell marker annotation by gene symbol
4. Epigenetic annotation & exosome-related genes annotation
5. GO cell biology function annotation.
6. KEGG pathway annotation
7. Enrichment analysis

Essentially, our challenge lies in the absence of comprehensive integration for the raw data. This deficit encompasses a lack of diverse tags, shared common information, and crucially, the biological mechanisms and knowledge. To predict uncertain relationships and tags, as well as to generate new cell samples and explore the entire cell state space, we can harness the power of deep generative models. As we know, increasing information leads to enhanced accuracy. The size of nodes, edges, and algorithm ability significantly impacts Graph Neural Networks (GNN). Given the wealth of information and knowledge, utilizing these models becomes essential.

In the realm of complex human diseases, intricate interplay among several molecules exhibits multi-level, cross-modal, and time-dependent characteristics, marked by interactions and connections. From a mathematical modeling perspective, the key challenge lies in quantifying these node relationships with specific biological significance, establishing suitable measurement approaches, and devising methods to meaningfully learn these relationships. Delineating whether particular relationships are genuine or possess direct causal links often necessitates extensive experimentation for validation, a process that can span years. Considering these limitations, the development of novel computational methods aimed at discovering and predicting potential relationships holds paramount importance. Such methods have the uncertainties associated with designing wet lab experiments and increase the overall efficiency of disease research endeavors.

In graph networks, the interpretation of the existence of node attributes is contingent upon the nature of connection between nodes and the corresponding connection weight. The significance of these attributes is closely tied to the specific type of connections leveraged between nodes and the associated weights assigned to these connections. However, the process of defining and comprehending the connection type of a relationship, along with the connection weight, through variation in node attributes is precisely handled what can be acquired through graph networks. This capability to learn and derive such insights is a fundamental reason, which argues why graph networks are being harnessed for data representation in the context of multi-omics. This approach is also especially beneficial over employing structured data frames for relationship representation. Graph networks excel at capturing complex relationships and nuances inherent in intricate data, making them preferred choice for modeling these interactions. Incorporating multi-omics data by using graph networks offer a two-fold advantage. It not only resolves the issue of over-sparse data frames representation, but also enable a more effective description of intricate interplay between nodes and their neighboring nodes. This efficacy stems from the fact that these interactions arise as a consequence of collaborative variations brough about by alterations in node attributes, relationship types, and their associated weights. Nevertheless, the attributes of nodes and their corresponding representations are dynamically interdependent and interconnected, adapting in tandem with modifications in interaction relationships. This approach enables a more accurate reflection of the relationships between nodes and their neighboring counterparts.

In this research, we employ the adaptive graph diffusion convolution method to capture the aforementioned dynamics information. This is comprised of two fundamental steps. First, we establish learnable parameters for each type of relationship. Subsequently, during the training phase, we engage in conditional adaptive graph diffusion convolution learning using cell subtypes as labels. This method fulfills our specific objective of generating cell samples of diverse subtypes. Taken together, these two steps constitute the central concept underlying our modeling strategy adopted in this research.

Furthermore, our method does not mandate the inclusion of exhaustive relational information as input to the model. Instead, we follow a two-step process. Initially, we derive vector representations of nodes by unsupervised learning. Thereafter, we delve into the interplay between node properties and edge types by assigning distinct learning weight coefficients to different categories of edge relationships. This enables us to capture the intricate relationships withing the data.

Subsequently, fine-tuning is undertaken using a small set of conditionally labeled data with supervision. This step partially tickles down the challenge of accurately modeling multi-omics heterogenous graph networks. This approach allows us to obtain an improved level of precision and insight within our modeling framework.

### 5. Improved part of our ACGDC model

The original GDC model significantly mitigates the overfitting problem associated with traditional GCN [75]. This is primarily due to the capability of GDC to counteract the noise inherent in edge relationships [76]. GDC obtains this by facilitating the smoothing of these edge relations. As a result, GDC is especially well-suited for capturing the intricate interplay between edges and nodes. This is obtained through the incorporation of node properties and the process of message passing. The fundamental intuition underlying GDC is akin to a smoothing process applied to the graph’s neighborhood, akin to the denoising function performed by Gaussian filters on images [77]. This approach substantiates beneficial for graph learning, as both the features and edges within actual graphs frequently exhibit noise. By incorporating this smoothing mechanism, GDC enhances the quality and reliability of the graph-based learning process.

However, it is crucial to note that GDC is primarily tailored for homogeneous graphs, which hinders its effectiveness when applied to heterogeneous graphs. Fortunately, the adaptive graph diffusion convolution (AGDC) approach presents a resolution for addressing the problems posed by heterogeneous graphs. AGDC achieves tis by incorporating learnable parameter for various edge types (multi-channels) [78]. For a comprehensive understanding of the specific details of AGDC, refer to the reference [78].

In this context, to generate cell samples based cell subtype labels, we draw inspiration from the Classifier-Guidance solution [79], rather than the Classifier-Free solution [80]. Given the substantial training costs associated with the diffusion model and the relatively manageable training demands of the classifier, we adopt a strategy that involves repurposing pre-trained unconditional diffusion model and leveraging classifiers to fine-tune the generation process. This approach generates the control generation process, termed the post-modified Classifier-Guidance scheme. Conversely, the prior training Classifier-Free scheme entails the incorporation of conditional signals into the training of the diffusion model to enhance generation outcomes. The most critical phase in generating the diffusion model is the generation process *p*(***x***_*t*−1_ ∣ ***x***_*t*_). In this context of generation with ***y*** serving as the input condition, the modification involves substituting *p*(***x***_*t*−1_ ∣ ***x***_*t*_) with *p*(***x***_*t*−1_ ∣ ***x***_*t*_, ***y***), signifying the incorporation of input y into the generation process.

To reuse the already trained unconditional generative model *p*(***x***_*t*−1_ ∣ ***x***_*t*_), we use Bayes’ theorem to obtain:

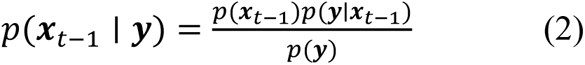

Complementing each term with the condition **x**_*t*_, we obtain:

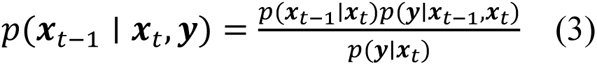

Note that in the forward process, **x**_*t*_ is obtained by ***x***_*t*−1_ by adding noise, and the noise will not contribute to the classification, so the addition of *x*_*t*_ will not have any gain for classification, so we have *p*(***y*** ∣ ***x***_*t*−1_, ***x***_*t*_) = *p*(***y*** ∣ ***x***_*t*−1_) and thus,

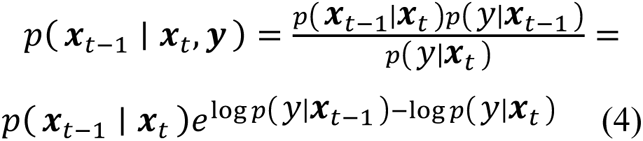

After the relevant derivation in the original paper, we only need to change the sampling of the generation process to

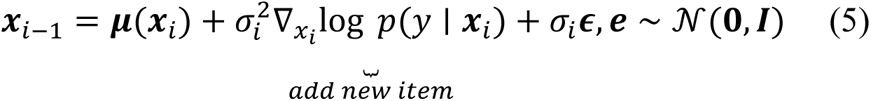

The whole framework of ACGDC is available at S7 in the supporting file.

## II. EXPERIMENTAL

### 1. Training parameters of ACGDC model

We conduct all experiments on a single NVIDIA RTX A5000 with 24 GB GPU memory. For our dataset, we utilize 3 ACGDC layers with transition matrix of GAT, hidden dimension 256, 6 attention heads, and residual linear connections.

### 2. Biological analysis and evaluation methods

To evaluate the effectiveness of the generated samples, we execute the experimental procedure outline below:

● Generate the samples by the trained ACGDC model.
● Perform several biological analyses between the generated samples and the real samples from different perspectives (enrichment analysis, cluster analysis, spatial distribution)
● Screen out the generated samples that can be used in the normal control group.
● Identify the subtype of cell for available samples from last step.

## III. RESULT

### 1. Analysis of raw gene expression profiles data

Before to initiating model training, an analysis of the original samples will be performed, encompassing the following aspects:

1. Heatmap Analysis at the Single Cell Level: evaluation of individual cell expression pattern by employing heatmap visualization.
2. Highly Expressed Gene Plot: Visualization of gene exhibiting high expression level across cells.
3. Cell Type Clustering Based on Expression: Categorizing cell types through clustering analysis of expression pattern.
4. t-SNE Plot of Brain Cell Types: Dimensionality reduction and visualization of brain cell types using t-SNE plots.
5. Age and Death Classification (DTHHRDY): Classification based on factors like age and death status.
6. KEGG and GO Term Analysis of the Marker Gene Set: Examination of enriched KEGG and GO terms associated with marker gene sets.

These analyses synergistically give insights into the characteristics and patterns within the actual sample data, laying the groundbreaking for subsequent model training.

### 2. Analysis of heatmap

To compare the gene expression difference among cells, a series of preprocessing steps are undertaken. Such steps are elaborated as:

1. Cells with Gene Count Filtering: Cells possessing fewer than 200 genes filtered out.
2. Genes with Cell Count Filtering: Genes occurring in less than 3 cells are filtered out.
3. Normalization: Each cell’s gene counts are normalized to ensure a total count of 1e6 genes per cell after normalization.
4. Mitochondrial Gene Removal: Mitochondrial genes are removed during the preprocessing of data.

After these steps, the dataset comprises 2642 cells and 54493 genes. In order to compare the gene expression variations, the observation variables of age and cell type are selected. Consequently, the resulting data is utilized to create heatmap figures depicted in Figure 8 and 9, illustrating the patterns and variations in gene expression across different cell types and ages.

In Figure 2, remarkable expression is observed for the genes ‘ENSG00000243649.8’ and ‘ENSG00000143476.17’ within the cell type of Brain - Anterior cingulate cortex (BA24) and Brain - Substantia nigra, respectively. Additionally, several other genes demonstrated expression biases among distinct cell subtypes.

**Figure 2.**
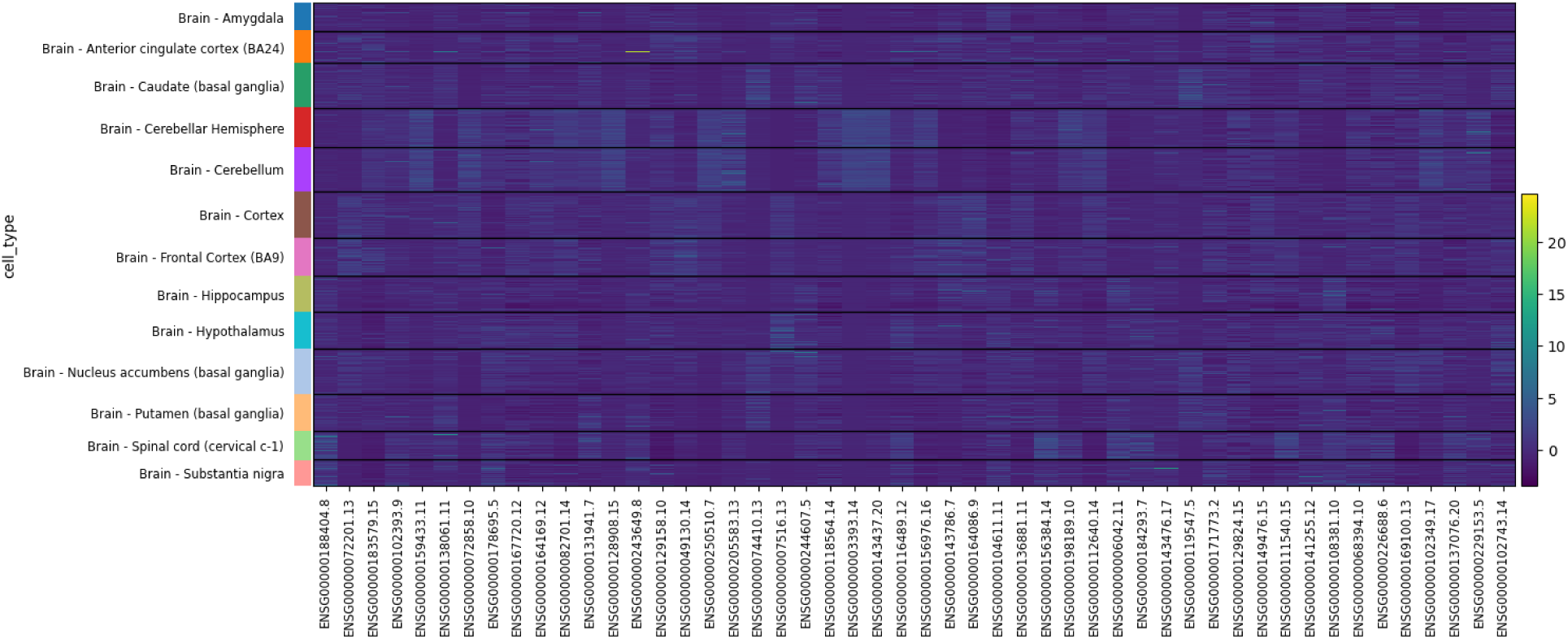
Differences in gene expression for different cell types. The color bar on the right indicates that the closer to yellow the higher the expression value, and vice versa. The different colors on the left side represent the range of intervals where different cell types are located, and the expression value of each gene in the same cell subtype is a distribution. The bottom represents the marker genes we selected, 50 in total.

In Figure 3, differential expression patterns are evident. Notably, the genes ‘ENSG00000243649.8’ and ‘ENSG00000102393.9’ showed significant expression within the age range of 60-69, while ‘ENSG00000229153.5’ exhibits high expression in the age range of 70-79. Moreover, many genes demonstrated expression biases across different age ranges.

**Figure 3.**
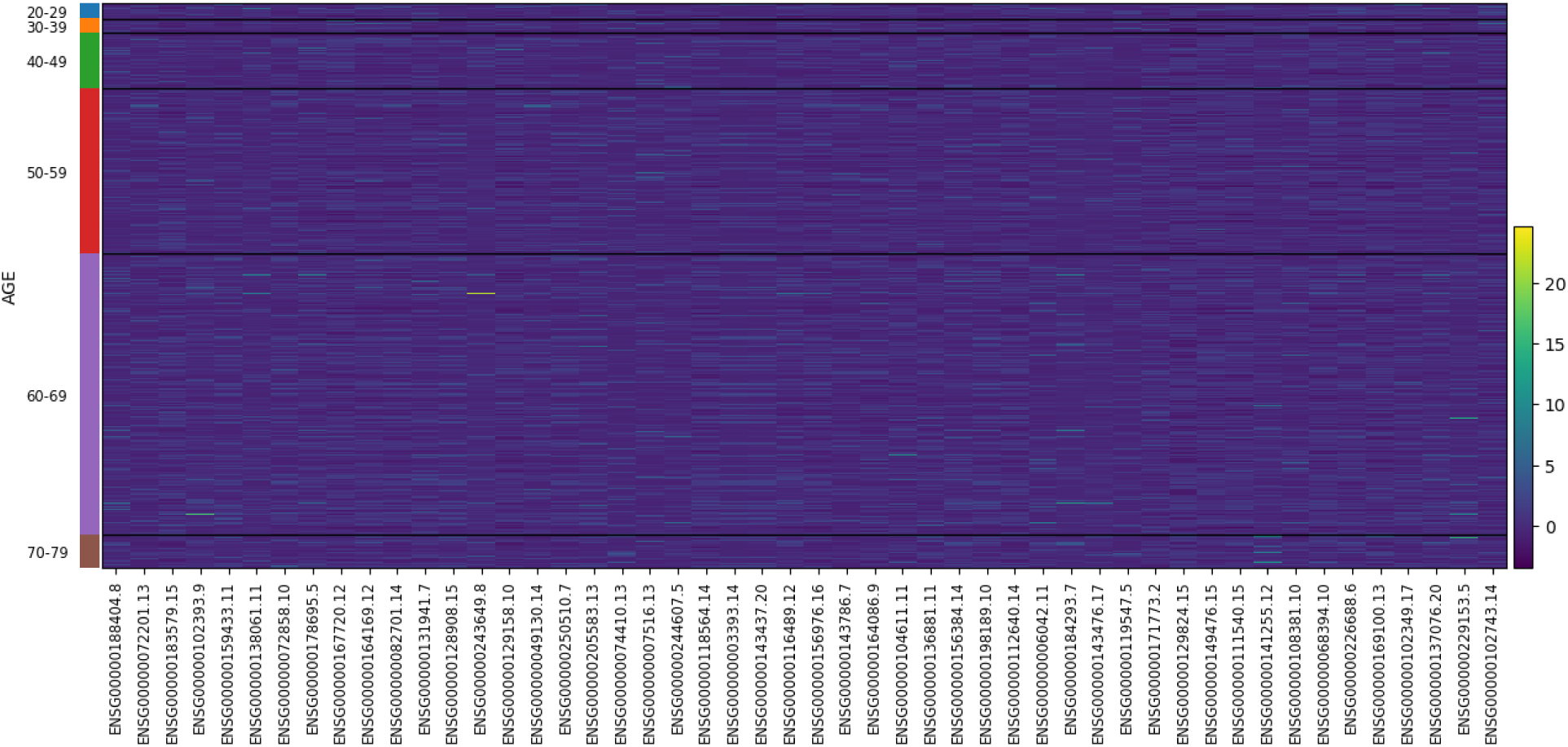
Differences in gene expression for different age ranges. The color bar on the right indicates that the closer to yellow the higher the expression value, and vice versa. The different colors on the left side represent the range of intervals where different age ranges are located, and the expression value of each gene in the same age range is a distribution. The bottom represents the marker genes we selected, 50 in total.

Figure 4 demonstrated a pronounced trend where exclusively the top three genes – MBP, GFAP, and PLP1 – display elevated levels of expression.

**Figure 4.**
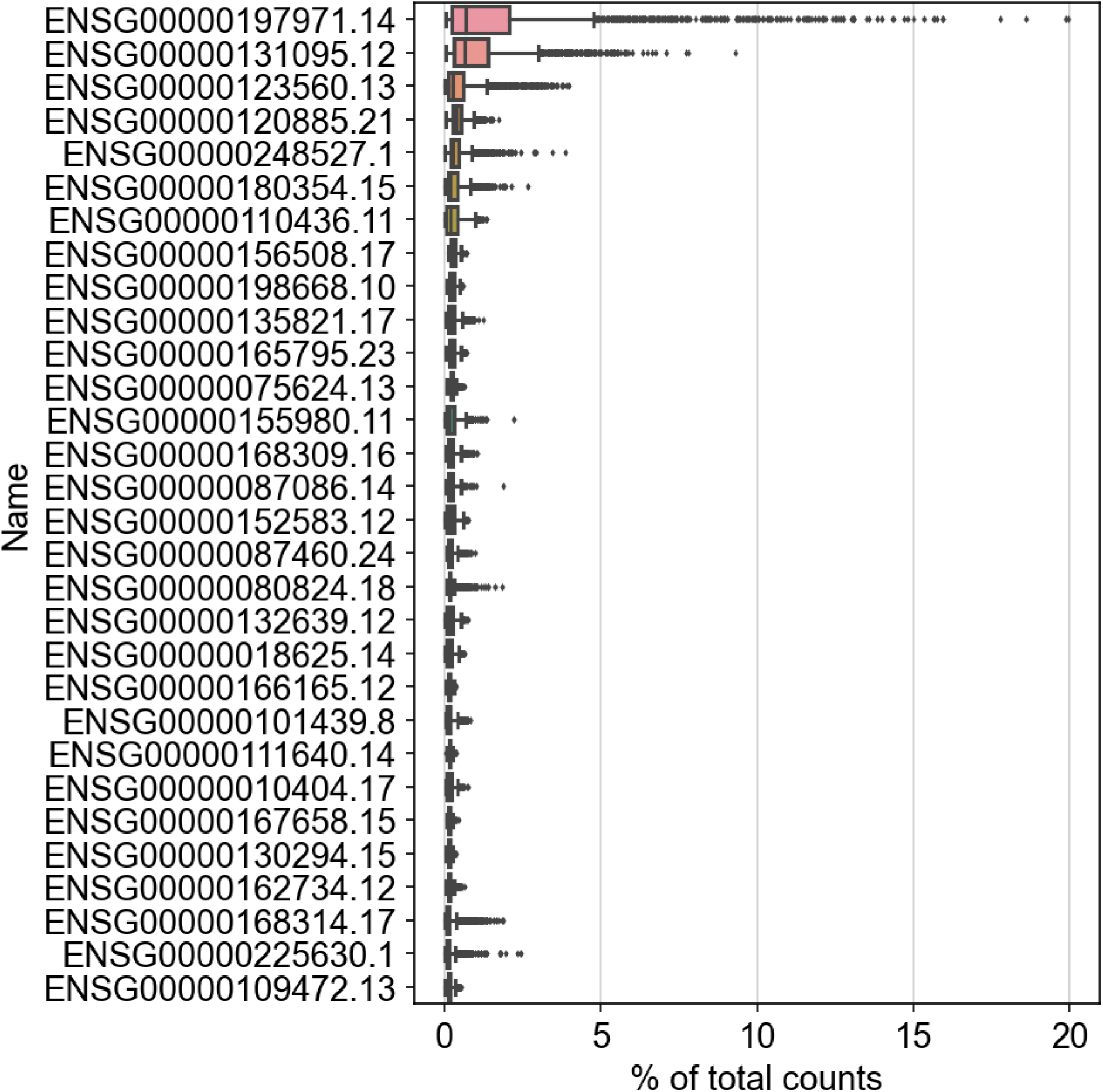
Top30 genes with the highest percentage of expression levels (excluding aberrantly expressed genes as well as mitochondrial genes).

### 3. Analysis of clustering and dimensionality reduction

We performed hierarchical clustering using cell types as the basis for single-cell gene expression matrices. In Figure 5, we observe the presence of 13 different cell types across all the samples. The hierarchical clustering output, based on the similarity of the gene expression vectors, indicated that a total of 5 subgroups were generated at the first level, 3 groups at the second level, 2 groups at the third level, 1 group at the fourth level, and finally, 1 group at the fifth level.

**Figure 5.**
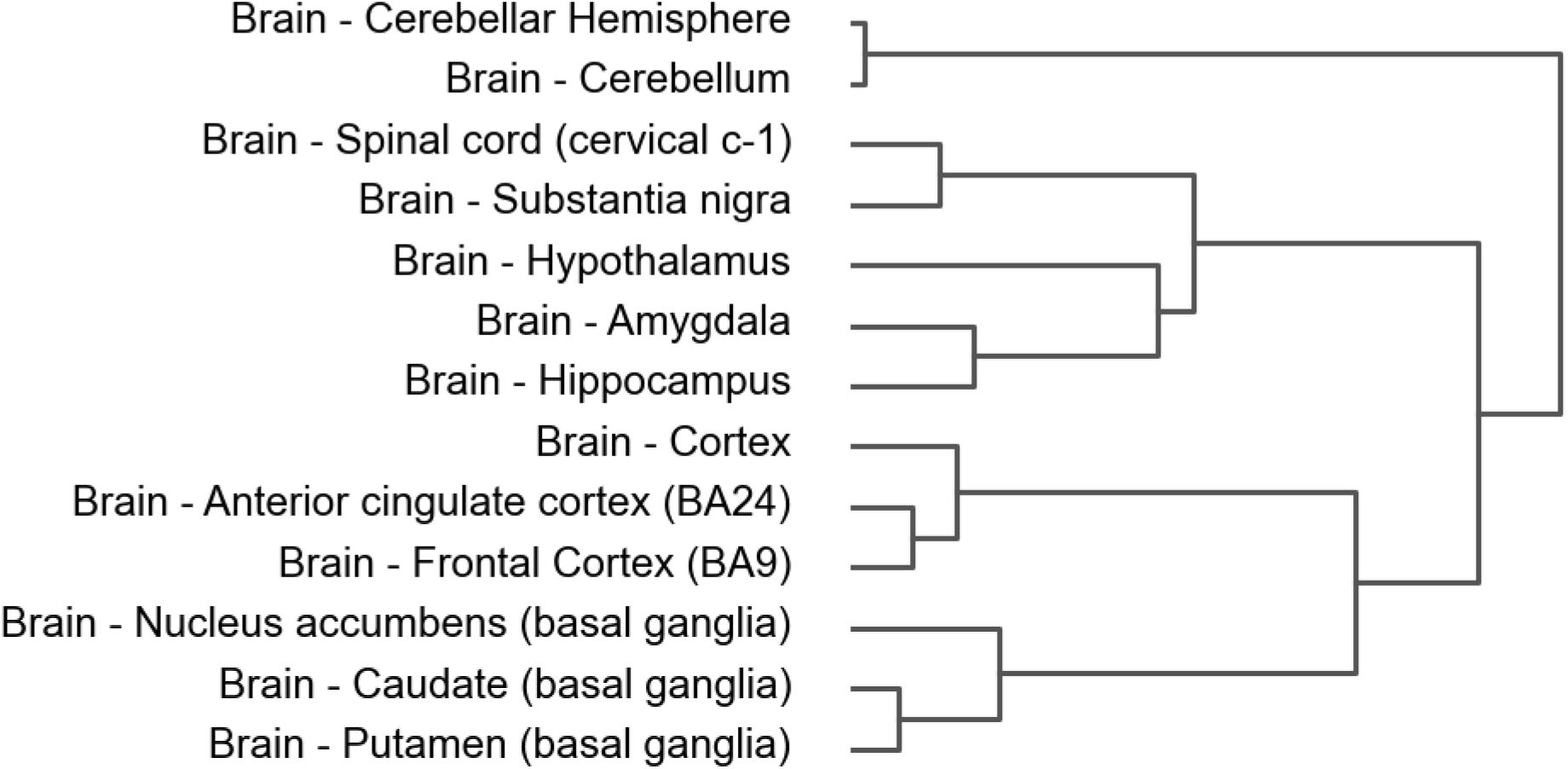
A hierarchical clustering map of brain cell types.

In Figure 6, our results observe the representation of 2642 cells in a 2D coordinate system obtained by downscaled gene dimensions employing t-SNE. The obtained plot displays that the 13 different cell types are clustered into three primary regions. Notably, cells represented by purple and red markers are closely grouped together and differentially distanced the other cell types. This clustering pattern demonstrates that the gene expression of these two cell types harbors a high degree of specificity.

**Figure 6.**
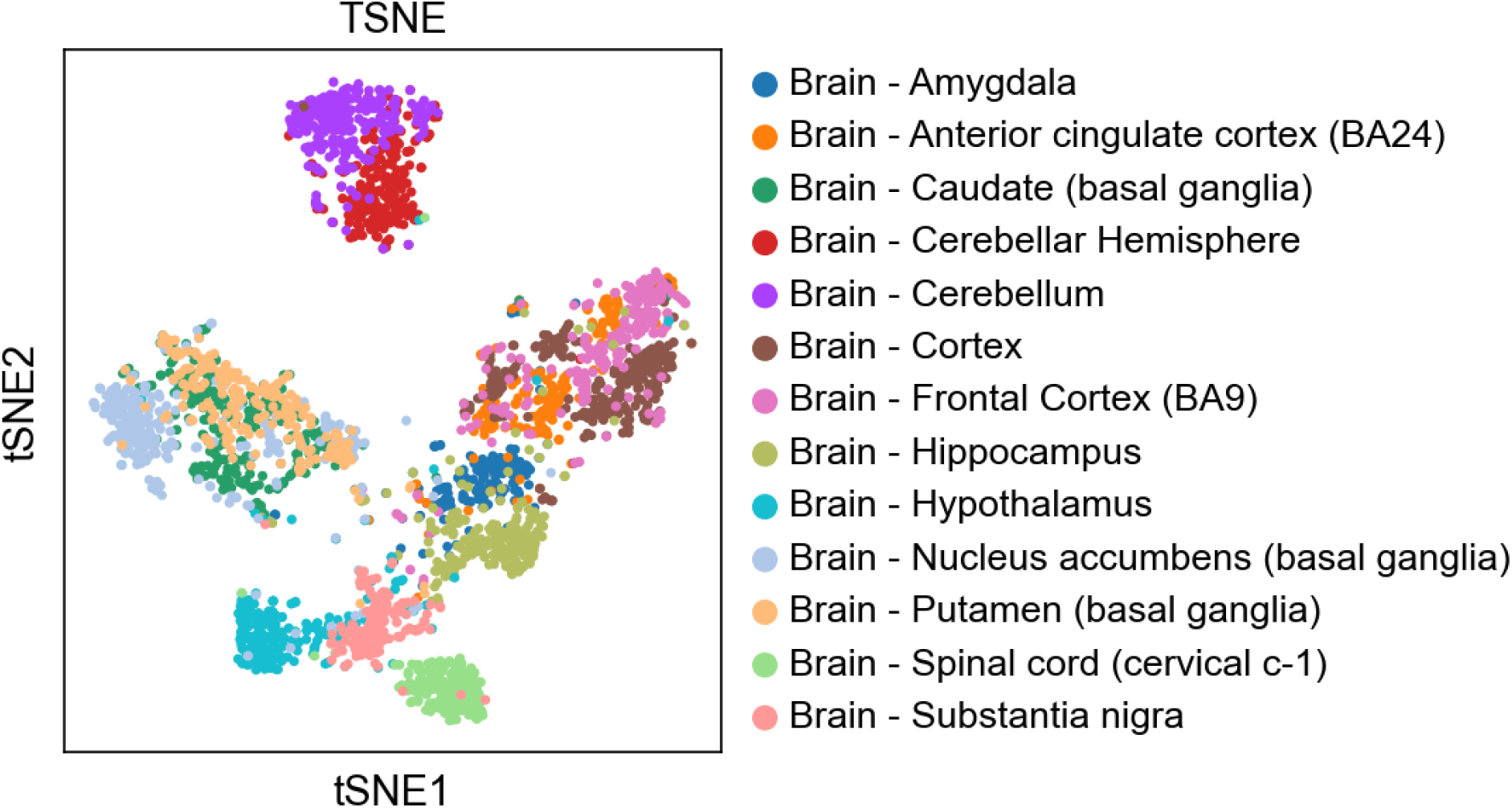
The t-SNE mapping of brain cell types.

Nevertheless, when considering age and death type as observed variables in Figure 7, specific gene expression patterns were not discernible. Instead, cells were distributed across various groups without remarkable trends. This observation points out that cell type, when treated as an observed variable, demonstrates greater specificity, and association with expression patterns of cells compared to age and death type.

**Figure 7.**
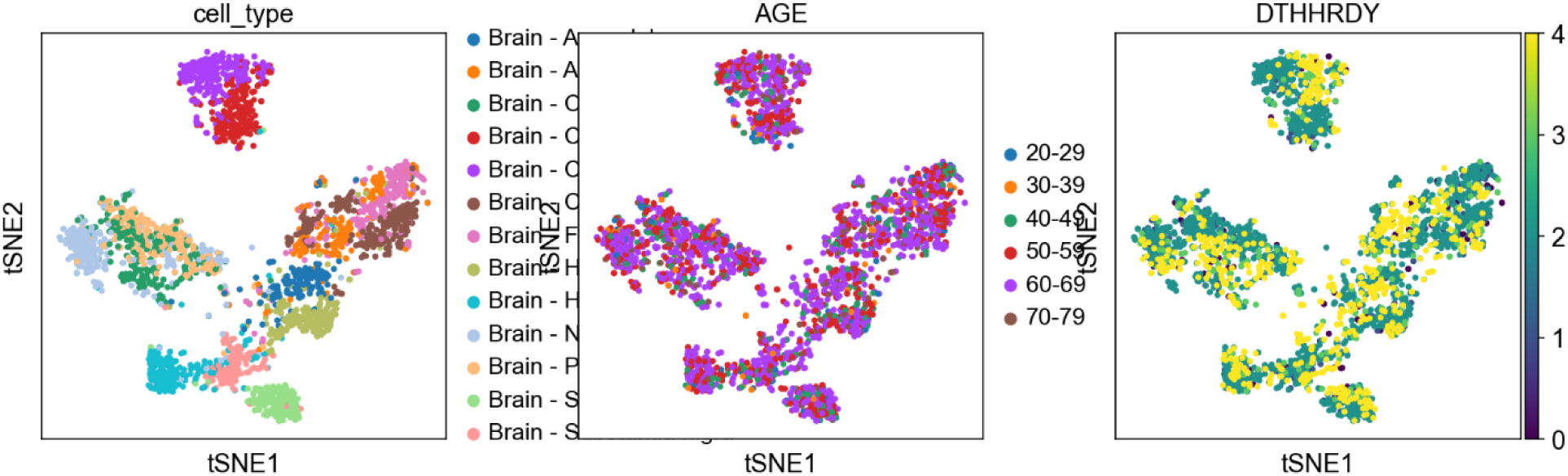
The t-SNE mapping of brain cell types, age and death classification (DTHHRDY).

### 4. Gene set enrichment analysis

Herein, we performed KEGG and GO term enrichment analysis of the 5658 genes that could be matched to the enriched term, as displayed in Figures 8 and 9. The results observed that these genes are predominantly enriched in pathways such as “Pathways in cancer”, “Olfactory transduction”, “MAPK and PI3K-Akt signaling pathway”, Human papillomavirus infection”, and “Olfactory Receptor Activity”.

**Figure 8.**
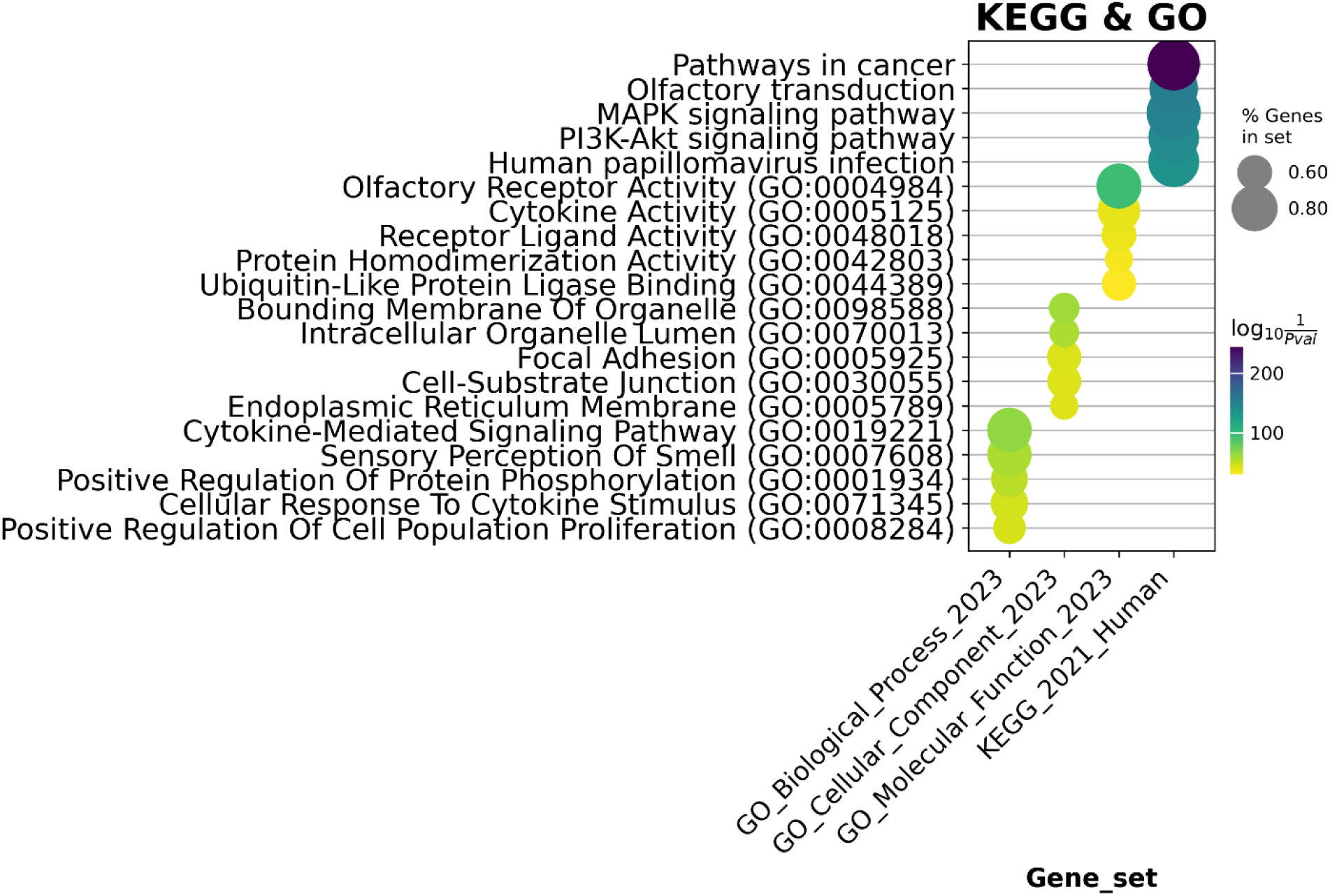
Top5 term for kegg&go gene set enrichment with dot plot

**Figure 9.**
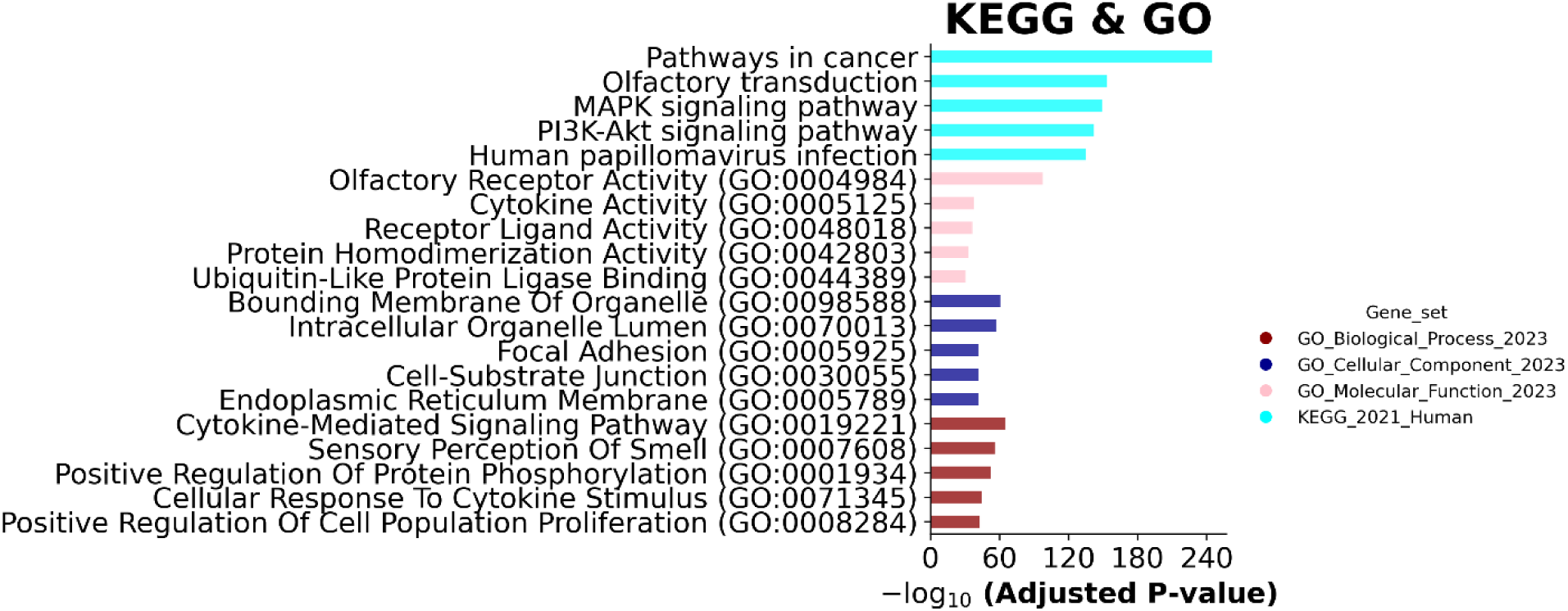
Top5 term for kegg&go gene set enrichment with bar plot

In the aforementioned section, we have performed an initial exploration and analyses of our original samples. We have focused on heatmap, clustering map, and highest gene expression plots illustrating the highest gene expression levels to understand gene expression patterns. In parallel, our aim is to generate similar but distinct samples from the original ones, here onwards, we will depict several generations results and compare the efficiency of the model.

### 5. Comparison of model performance

Since single-cell gene expression profiles typically exist in the form of 2D matrices, while clinical data as well as other annotated information are available in pairs. Algorithms designed for handling single-cell multi-omics data primarily include non-sequential algorithms. For instance, there exist VAE-based approach and the GAN-based approach, which treat a single cell/gene expression profile as a sample. Additionally, graph network approaches such as GDC and the F2GNN approach are employed, where the latter treats a first-order sub-network containing nodes as samples [81].

Our proposed method, ACGDC, is an improvement of GDC, an approach initially designed for homogenous graphs. Nevertheless, ACGDC addresses the limitations posed by diverse node and edge types. To obtain this, we embed note type information into their attributes, effectively converting various node types into a unified homogenous representation. In parallel, we encode distinct edge types and seamlessly append them into the attributes of edges. This adaptation empowers ACGDC to effectively operate on graphs with heterogenous node and edge types, enabling a more comprehensive and versatile analyses of intricate data structures. Nevertheless, considering all edges as homogeneous graphs and assigning them equal weights, an essential assumption is made that may not prove true in real-world experimentations. Realizing this challenge, our method presents a more nuanced solution. We assign distinct parameters to the first-order connected edges of each node, enabling the machine learning process to learn varying weights that are more reflective of the actual relationships. Conclusively, this method not only improves the interpretability and modeling accuracy but also aligns better with the unique objectives of the target task. Table 1 illustrates an overview of the performance of major single-cell generation models, focusing on matrices like validation rate, average similarity, and novelty.

**Table 1.** Comparison with different models for generation performance of single cell sample.

| Model | Validation Rate | Average Similarity | Novelty |
| --- | --- | --- | --- |
| VAE-normal | 0.45 | 0.4476 | 0.54 |
| WGAN-gp-normal | 0.34 | 0.3369 | 0.44 |
| GDC | 0.55 | 0.5512 | 0.65 |
| F2GNN | 0.50 | 0.4991 | 0.58 |
| ACGDC | 0.59 | 0.5884 | 0.68 |

Similarity: similarity to a nearest neighbor is an expression-valued Euclidean distance. Validation Rate: the ratio of maximum similarity >0.3 for all the generated samples.

Average Similarity: average score of similarity of the generated samples in the same subtype. Novelty=Number of generated samples not in training set / Number of unique and valid generated samples

In Table 1, the “-normal” designation indicates the use of min-max normalization and inverse normalization on both the training gene expression matrix data and the generated new samples. These findings observed that the WGAN based approach outperforms the VAE method across all three metrics. Among the three graph networks, F2GNN shows better results compared to GDC in terms of all three metrics. Nonetheless, ACGDC surpasses both GDC and F2GNN in performance as it assigns distinct parameters to each edge type for weight learning. In the subsequent analyses, our study will focus on comparing the samples generated by ACGDC model with the original training samples.

### 6. Biological analysis of generated cell gene expression profile

To assess the differences between the generated samples and the original samples, the preliminary step involved the generation of new samples employing ACGDC model. Consequently, a meaningful analysis was performed, constraining the newly generated samples with the original samples. This analysis surpassed several aspects including the ranking of differentially expressed genes, hierarchical clustering based on cell type, dimensionality reduction using PCA, and clustering visualization through UMAP.

At first, we generate a set of 30,000 samples employing the trained model. Thereafter, we subject these samples to preliminary filtering based on specific gene and cell criteria. Following the filtering, we proceed to normalize the gene expression by formula (6), and then execute the K-NearestNeighbor (KNN) algorithm to classify these cells based on their gene expression patterns. The term “validation sample” pertains to a sample that aligns with a cluster of specific cell subtypes.

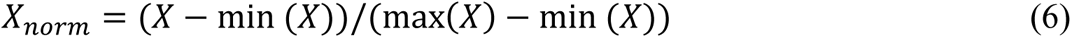

Figure 10 demonstrates the gene distribution of the top 30 genes, where A indicates the original sample and B represents the generated sample employing the ACGDC model. It is evident that the gene distribution ranking generated by the ACGDC model closely resembles that of the original sample. The top 7 genes are solely consistent, and a total of 24 genes are common among the top 30 genes, accounting for 80%. However, there is a remarkable difference in the numerical distribution, primarily due to the utilization of maximum-minimum normalization algorithm, resulting in value comparison in the generated samples, it does not impact the overall distribution pattern.

**Figure 10.**
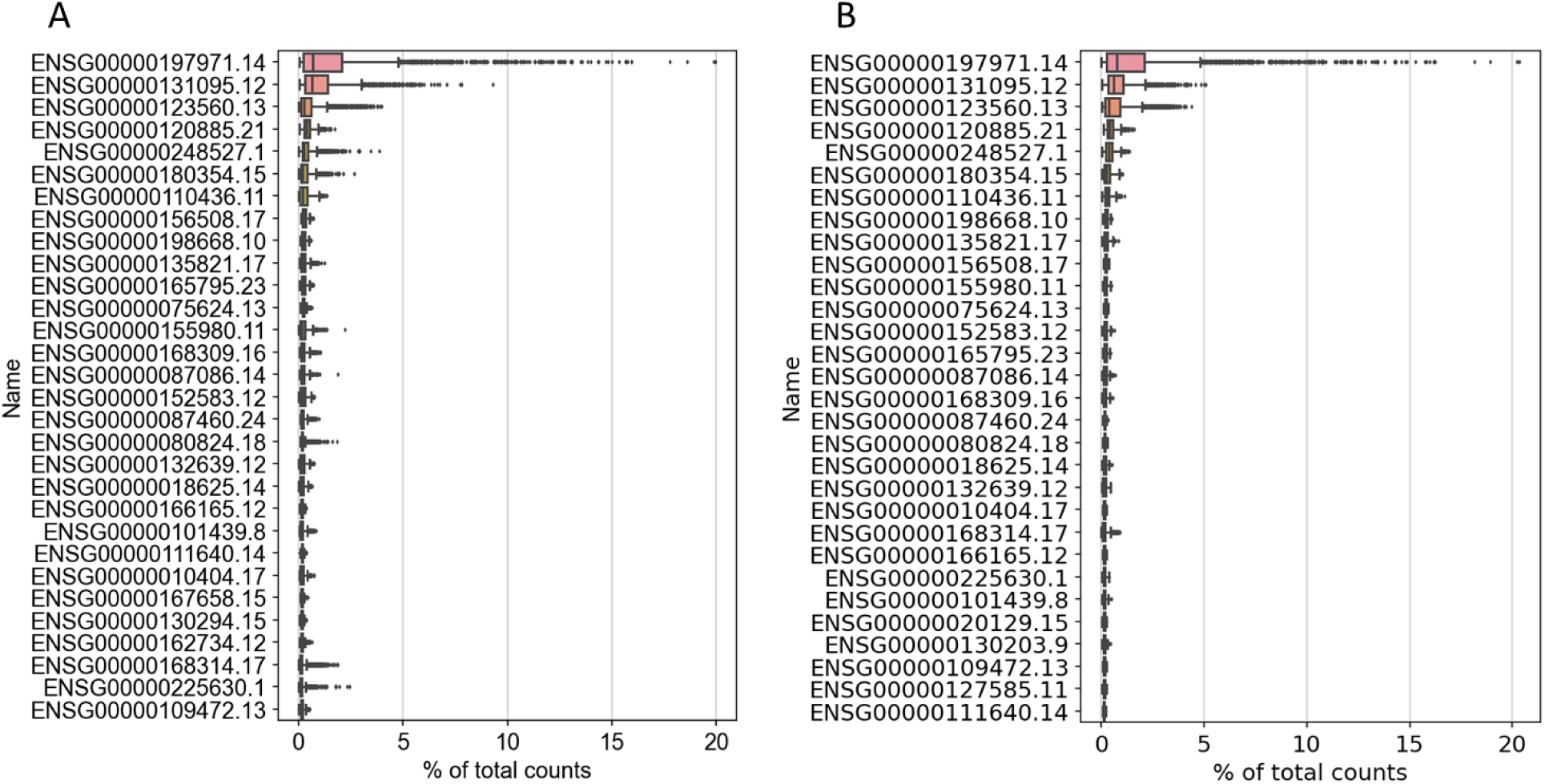
Top30 gene rank (% of total counts). A means original sample, B means generated sample.

For comparison with the original sample space, Figure 11-13 demonstrates the clustering results for the generated samples.

**Figure 11.**
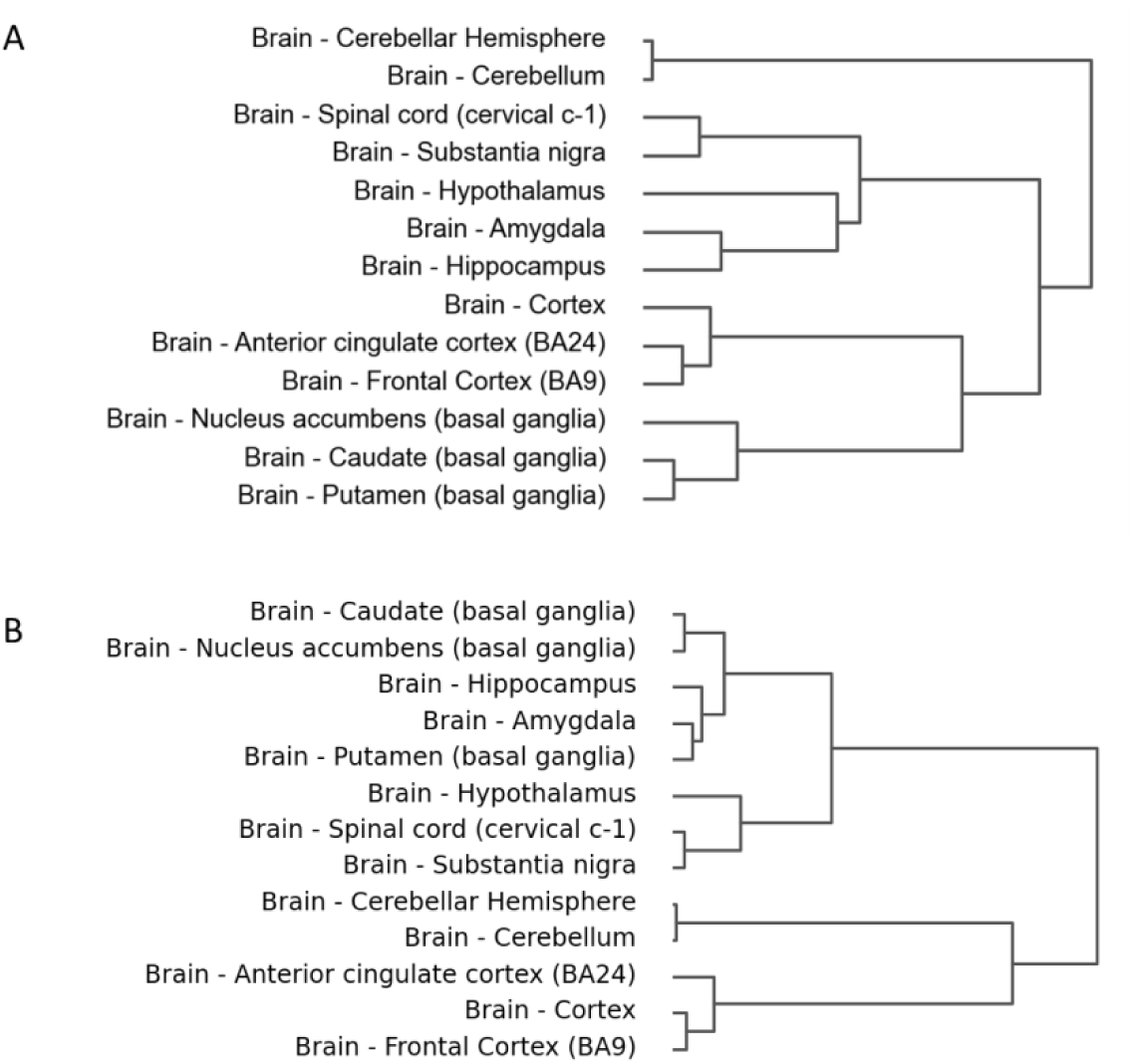
Hierarchical clustering based on gene expression profiles of cells and cell subclasses. A means original sample, B means generated sample.

In Fig 11 A and B, we observe that Brain - Cerebellar Hemisphere and Brain – Cerebellum, as well as Brain - Spinal cord (cervical c-1) and Brain - Substantia nigra are clustered to the same class in the first-order clustering outcomes. In parallel, Brain – Hypothalamus, Brain – Amygdala, Brain – Hippocampus, and Brain - Anterior cingulate cortex (BA24), Brain - Frontal Cortex (BA9), Brain – Cortex, and Brain - Putamen (basal ganglia), Brain - Nucleus accumbens (basal ganglia), Brain - Caudate (basal ganglia) are all clustered into the same class in the second-order clustering results.

Figure 12 indicates that there is a clear similarity in the distribution between the generated samples and the original samples, especially evident in the case of the cell subtypes Brain - Cerebellar Hemisphere and Brain - Cerebellum.

**Figure 12.**
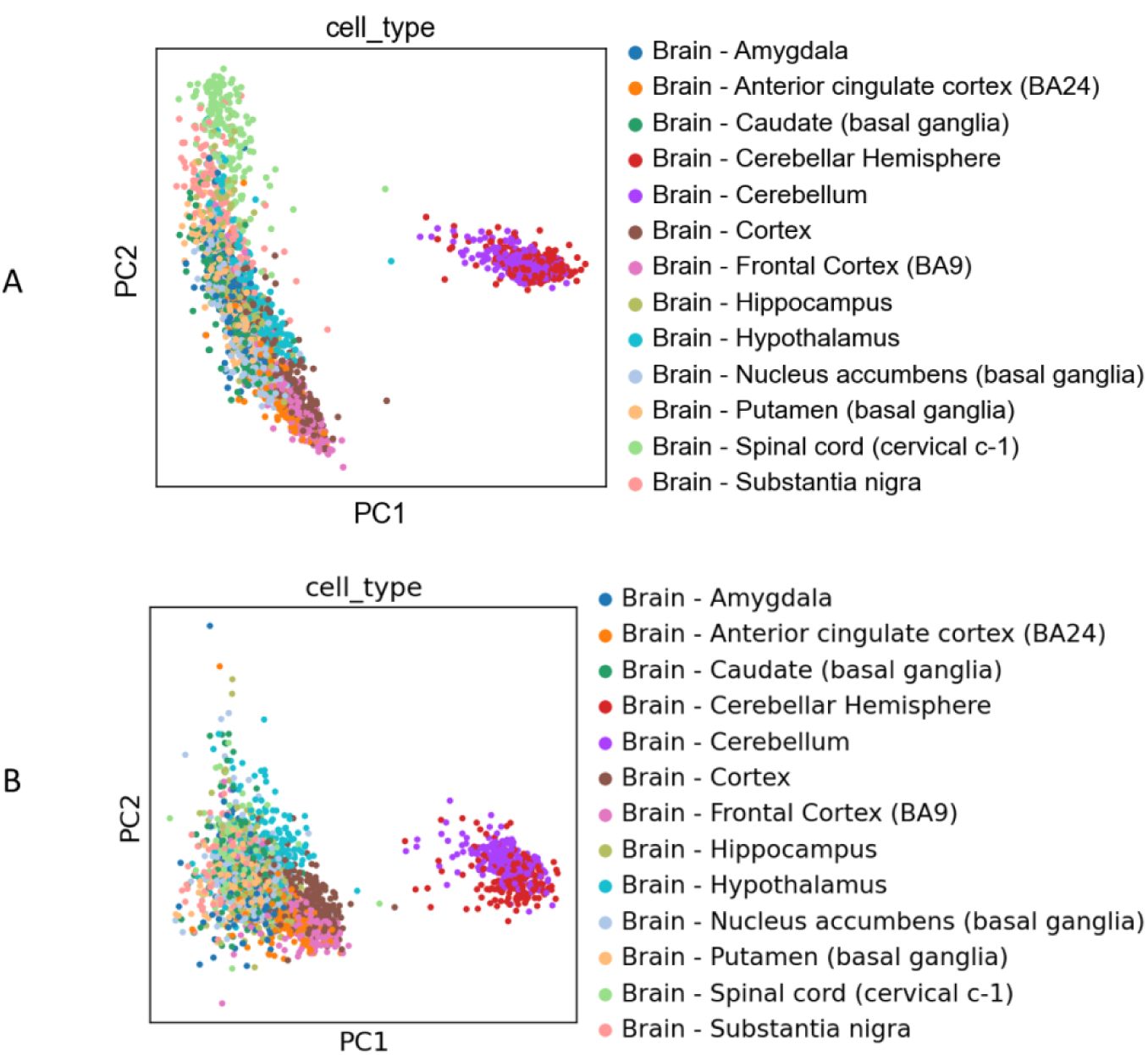
PCA clustering based on gene expression profiles of cells and cell subclasses. A means original sample, B means generated sample.

We perform Louvain algorithm for clustering before applying UMAP mapping [82, 83].

Figure 13 illustrates that the distribution of generated samples and original samples are generally similar, since the generated sample can be clustered to more classes.

**Figure 13.**
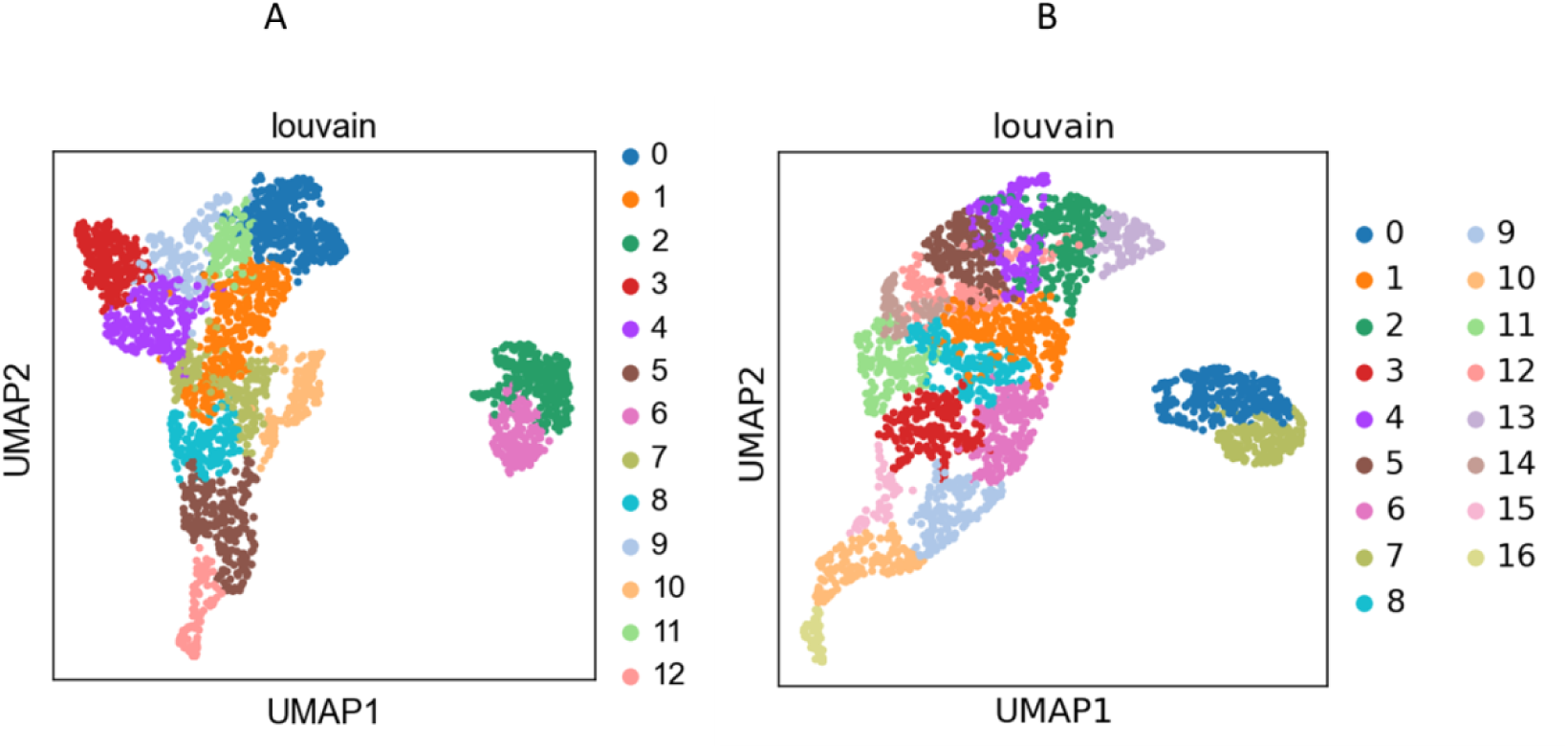
UMAP-Louvain clustering based on gene expression profiles of cells and cell subclasses. A means original sample, B means generated sample.

### 7. Comparison with experiment samples and availability screening for different cell subtypes

In addition, we calculate the average similarity across all cells of the predicted category as depicted in Table 2. Consequently, we treat samples with similarity greater than 0.5 as validated samples, as indicated in Table 3. The similarity is derived from the probability score predicted by KNN classifier.

**Table 2.** Descriptive statistics of screened samples belong to a cluster center of the cell subtype.

| Cell Subtype | Average Similarity | Count | Max | Min | Variance |
| --- | --- | --- | --- | --- | --- |
| Brain - Amygdala | 0.3609 | 68 | 0.5385 | 0.2308 | 0.0061 |
| Brain - Anterior cingulate cortex (BA24) | 0.5171 | 18 | 0.6923 | 0.3846 | 0.0082 |
| Brain - Caudate (basal ganglia) | 0.4745 | 71 | 0.6154 | 0.3077 | 0.0095 |
| Brain - Cerebellar Hemisphere | 0.6346 | 72 | 1.0000 | 0.5385 | 0.0080 |
| Brain - Cerebellum | 0.7706 | 220 | 1.0000 | 0.3077 | 0.0136 |
| Brain - Cortex | 0.5780 | 173 | 0.8462 | 0.1538 | 0.0235 |
| Brain - Frontal Cortex (BA9) | 0.4348 | 23 | 0.5385 | 0.3077 | 0.0052 |
| Brain - Hippocampus | 0.3454 | 53 | 0.4615 | 0.2308 | 0.0051 |
| Brain - Hypothalamus | 0.3504 | 9 | 0.4615 | 0.3077 | 0.0031 |
| Brain - Nucleus accumbens (basal ganglia) | 0.3846 | 86 | 0.6923 | 0.2308 | 0.0116 |
| Brain - Putamen (basal ganglia) | 0.4054 | 111 | 0.7692 | 0.2308 | 0.0119 |
| Brain - Spinal cord (cervical c-1) | 0.9196 | 133 | 1.0000 | 0.3846 | 0.0216 |
| Brain - Substantia nigra | 0.5304 | 76 | 0.7692 | 0.2308 | 0.0193 |
| **Total** | **0.5884** | **1113** | **1.0000** | **0.1538** | **0.0487** |

**Table 3.** Descriptive statistics of screened samples belonging to the cluster center of a cell subtype and similarity >=0.5.

| Cell Subtype | Average Similarity | Count | Max | Min | Variance |
| --- | --- | --- | --- | --- | --- |
| Brain – Amygdala | 0.5385 | 2 | 0.5385 | 0.5385 | 0.0000 |
| Brain - Anterior cingulate cortex (BA24) | 0.5846 | 10 | 0.6923 | 0.5385 | 0.0029 |
| Brain - Caudate (basal ganglia) | 0.5692 | 30 | 0.6154 | 0.5385 | 0.0015 |
| Brain - Cerebellar Hemisphere | 0.6346 | 72 | 1.0000 | 0.5385 | 0.0080 |
| Brain - Cerebellum | 0.7727 | 219 | 1.0000 | 0.5385 | 0.0126 |
| Brain – Cortex | 0.6560 | 125 | 0.8462 | 0.5385 | 0.0074 |
| Brain - Frontal Cortex (BA9) | 0.5385 | 4 | 0.5385 | 0.5385 | 0.0000 |
| Brain - Hippocampus | — | — | — | — | — |
| Brain - Hypothalamus | — | — | — | — | — |
| Brain - Nucleus accumbens (basal ganglia) | 0.5799 | 13 | 0.6923 | 0.5385 | 0.0026 |
| Brain - Putamen (basal ganglia) | 0.5662 | 25 | 0.7692 | 0.5385 | 0.0029 |
| Brain - Spinal cord (cervical c-1) | 0.9502 | 125 | 1.0000 | 0.5385 | 0.0073 |
| Brain - Substantia nigra | 0.6423 | 40 | 0.7692 | 0.5385 | 0.0062 |
| **total** | **0.7357** | **665** | **1.0000** | **0.5385** | **0.0238** |

— Means not exist.

As shown in Table 2, the findings revealed that ACGDC model has generated a total of 1113 validated samples, encompassing all the cell subtypes. The cumulative similarity score is 0.5884, indicating that 1113 samples can effectively replace 58.84% of the experimental samples. Furthermore, new samples from Brain - Spinal cord (cervical c-1), and Brain – Cerebellum, Brain - Cerebellar Hemisphere indicate exceptional quality based on their high average similarity, and large count, and low variance.

Finally, we screened and selected new samples with similarity of equal or more than 0.5, which can be used as negative samples for the control group, as outlined in Table 3. The details for each new sample, classified by different cell subtypes, are available in the supporting document.

As shown in Table 3, the ACGDC model has generated 665 samples, involving 11 different cell subtypes. The total similarity is measured at 0.7357, indicating that these 665 samples can effectively replace 73.57% of the experimental samples. Nevertheless, it’s noticeable that the cell subtypes “Brain – Hippocampus” and “Brain – Hypothalamus” have been not generated with meeting the condition of similarity equal to or greater than 0.5.

## IV. DISCUSSION

In this research, we have proposed ACGDC model for novel samples generation. Through this method, we successfully identified a total of 1113 validated samples, encompassing all the cell subtypes. Among these, we generated 665 high-quality samples, representing 11 distinct cell subtypes. These generated samples closely resembled the training set, establishing them as appropriate candidates for use as control samples. Our methods showed that top 30 genes’ ranking and the clustering outcomes are highly reliable and congruent in both gene expression profiles and the distribution of cell subtypes. Furthermore, our ACGDC model preferentially integrates several forms of annotated information including genomic and clinical data, as well as known cell types. It outperforms well-known generation models in terms of validation rate, average similarity, and novelty. This integrated approach offers an innovative solution to overcome the limitations associated with integrating single-cell multi-omics data. Nevertheless, it is evident that despite our model’s superior performance on numerous performance matrices, it encounters difficulty generating high-quality samples for two specific cell subtypes. Additionally, there is a limited fraction of samples generated for certain other cell subclasses. This limitation can be attributed to the lack of sample data within these categories. To address this limitation, future endeavors could engage research efforts by considering training the model with more diverse sample datasets. Certainly, this approach would not merely support the generation of samples across a broader spectrum of categories but also enhance the overall quality of the generated samples.

## V. CONCLUSION

Astrocytes play a pivotal role in the study of brain diseases such as GBM, and AD, as well as in the study of brain mechanisms like sleep, learning and memory. Notably, there is no efficient drug available in the market for GBM, making targeted therapy a beacon of hope for this disease. Moreover, obtaining clinical samples of astrocytes, whether normal control or disease samples, encompassing several subtypes and their transitional states, poses considerable challenges. Therefore, the development of methods for generating more closely similar samples from the limited available samples, and even venturing into uncharted territories biological features represented by these samples, holds significant potential to alleviate the bottleneck created by the scarcity of astrocyte samples in research.

In the realm of computational biology and drug discovery, contemporary deep generative models have made breakthroughs. Deep learning methods including GAN, GCN, GAT, GDC, and Transformer have been introduced into the single-cell research domain. They are leveraged to facilitate sample space learning under specific conditions and for distinct objective. This integration has remarkably contributed to the generation of novel samples, including single-cell types and modeling of single-cell state trajectories.

This paper demonstrates a feasible and efficient model for GNNs. First, we refine Adaptive Graph Diffusion Convolution Networks (AGDCNs) by replacing the graph convolution operator with a more efficient graph diffusion method in each GNN layer. Subsequently, we extend this graph diffusion concept to introduce the Adaptive Conditional Graph Diffusion Model (ACGDC), which incorporates conditional sampling from the AGDCNs. We evaluate ACGDC and other popular GNNs on node classification and conditional generation. The experimental findings reveal that ACGDC obtained significant superiority over popular generation models like VAE and GAN, as well as state-of-the-art GNNs such as F2GNN and GDC.

In summary, our approach entangles the integration of single-cell multi-omics data through a heterogeneous graph network. We employ message passing and graph diffusion convolution techniques to assign different weights to various edge types within the network. This innovative method, known as Adaptive Conditional Graph Diffusion Convolution (ACGDC) model, is designed to pre-train on a large-scale dataset of astrocyte samples and subsequently generate new control samples in downstream conditional generation tasks. The primary goal is to alleviate the scarcity of normal control astrocyte samples, particularly in the highly complicated field of brain disease research. Moreover, this approach aims to facilitate research into sleep and memory learning mechanisms while promoting advancements in the diverse domain of astrocyte-related research, including neurological diseases such as GBM and AD, as well as investigating into brain mechanisms.

Despite promising application of ACGDC model in brain disease research, it has certain limitations. One common issue, which is applicable to deep Graph Neural Networks (GNNs) like ACGDC, is the challenge of implementing node- or layer-neighborhood sampling techniques effectively. Moreover, the volume of available data plays a crucial role in determining the model’s effectiveness. To this end, future research will need to address several aspects, including:

**Scalability:** Developing methods to apply node- or layer-neighborhood sampling effectively in very deep GNNs is a challenge that requires further exploration.

**Data Volume:** Increasing the model’s performance would highly benefit by the integration of larger and diverse datasets. This include gathering more diverse samples and generating samples from a broader range of subtypes.

**Sample Quality:** Continual efforts should be made to improve the quality of generated samples, ensuring they closely resemble real-world data.

These limitations highlight avenues for future research and improvement in the application of ACGDC and similar moles in the field of computational biology and beyond.

## Author contributions

J.M conceived the idea and wrote the article, developed the codes, and prepared the figures. A.Z. aided to draw some figures, editing the language expression, K.N. provided the financial support, J.W. provided some advice on writing skills. All the authors discussed the results and commented on the manuscript.

## Acknowledgements

This work was supported by Yonsei University graduate school “Integrative Biotechnology & Translational Medicine” and the Establishment and demonstration of bio-material data platform, P0014714.

## ASSOCIATED CONTENT

Supporting file is available at https://doi.org/10.5281/zenodo.8358553: additional experimental details, including generative samples for specified subtype of cell and original samples and annotated information.

The GTEx website: https://gtexportal.org/home/downloads/adult-gtex#bulk_tissue_expression.

